# Accumulation of Extracellular GABA, Impaired GABAergic Neurotransmission and 4-Phenylbutyrate Rescue in Mice of *SLC6A1* Variant-Mediated Disorders

**DOI:** 10.64898/2026.01.20.700615

**Authors:** Kirill Zavalin, Karishma Randhave, Marshall Biven, Wangzhen Shen, Jing-Qiong Kang

## Abstract

Mutations in *SLC6A1* encoding GABA transporter 1 are a leading monogenic cause of developmental and epileptic encephalopathies, severe neurodevelopmental disorders lacking effective treatments. We previously demonstrated that 4-phenylbutyrate restored molecular and functional deficits, and reduced seizures in a *Slc6a1* loss-of-function mouse, motivating a promising ongoing clinical trial. Here, we show this mouse exhibits accumulation of extracellular GABA, impaired neurotransmission, and reduced GABA uptake; and demonstrate that 4-phenylbutyrate rescues these abnormalities.

## Main Text

Pathogenic variants of *SLC6A1* encoding γ-aminobutyric acid (GABA) transporter 1 (GAT-1) are an important and increasingly recognized cause of developmental and epileptic encephalopathies (DEEs), debilitating neurologic disorders marked by early-onset seizures and severe neurodevelopmental deficits^1-6^. A high demand exists for developing novel therapeutics for *SLC6A1*-DEEs, as treatment options are limited to anti-convulsive medicines that do not stop all seizures and rarely help with comorbidities^2,3^.

*SLC6A1*-DEEs are thought to arise from pathologic changes in developmental signaling and inhibitory neurotransmission mediated by GABA, including synaptic and tonic inhibition through ionotropic GABA_A_ receptors, and modulation through metabotropic GABA_B_ receptors^7-10^. GAT-1 mediates the majority of GABA uptake after synaptic release, affecting the strength and duration of inhibition, and recycling GABA to GABAergic neurons^7-10^. When GAT-1 activity is impaired in GAT-1 knockout animals and with GAT-1 blockers, deficient GABA clearance impacts inhibitory neurotransmission, particularly GABA_A_ tonic and large synaptic currents^11-16^.

Our lab has extensively studied the patho-mechanisms of *SLC6A1*-DEEs and identified 4-phenylbutyrate (PBA) as a promising therapeutic. GAT-1 function and expression are decreased in most *SLC6A1* disease variants, which all exhibit a pronounced pathology in intracellular trafficking^17-19^. This is exemplified by GAT-1(S295L) complete loss-of-function variant, which causes seizures and neurodevelopmental comorbidities in knock-in *Slc6a1*^+/S295L^ mice similar to the patient^2,4,18,20^. Known as a chemical chaperone and histone deacetylase inhibitor^21^, PBA restored these pathologies for multiple *SLC6A1* variants *in vitro*^18,22^; and *in vivo* in *Slc6a1*^+/S295L^ mice, reducing seizure burden by over 70%^18^. Similarly, PBA helped in *Gabrg2*^+/Q390X^ Dravet-Syndrome mice^23^. These preclinical findings prompted a clinical trial in which 70% of patients with *SLC6A1* variants experienced seizure reduction or resolution (in submission)^24^. However, up to date, there is no study characterizing the GABAergic neurotransmission nor extracellular GABA level *in vivo* for *SLC6A1*-DEEs. Here, we interrogated disease mechanisms by assessing GABA uptake, GABA levels, and GABAergic transmission in *Slc6a1*(S295L) mice, and evaluating effects of PBA treatment.

We started by evaluating GABA uptake in *Slc6a1*^+/S295L^ mice and wildtype *Slc6a1*^+/+^ littermates by *ex vivo* H^3^-GABA uptake assay in gliosomes and synaptosomes extracted from brains at 2-4 months old. Consistent with our previously reported findings^19^, uptake was reduced in both *Slc6a1*^+/S295L^ synaptosomes (Figure 1A, 3641 vs 7255 CPM WT, ***P<.001) and gliosomes (Figure 1B; 2386 vs 4371 CPM WT, **P<.01). In synaptosomes, this deficit persisted when compensatory uptake was blocked with GAT-3 blocker SNAP5114 (Figure 1A; 2825 vs 5440 CPM WT, ***P<.001) but was normalized by GAT-1 blocker CL-966 (Figure 1A; 2098 vs 2042 CPM WT), indicating that deficiency was specifically attributable to loss of GAT-1 function. Results were less clear in *Slc6a1*^+/S295L^ gliosomes, where decreased uptake persisted with CL-966 (Figure 1B; 1023 vs 1957 CPM WT, ***P<.001) and SNAP5114 (Figure 1B; 1698 vs 3801 CPM WT, **P<.01).

**Figure 1.**
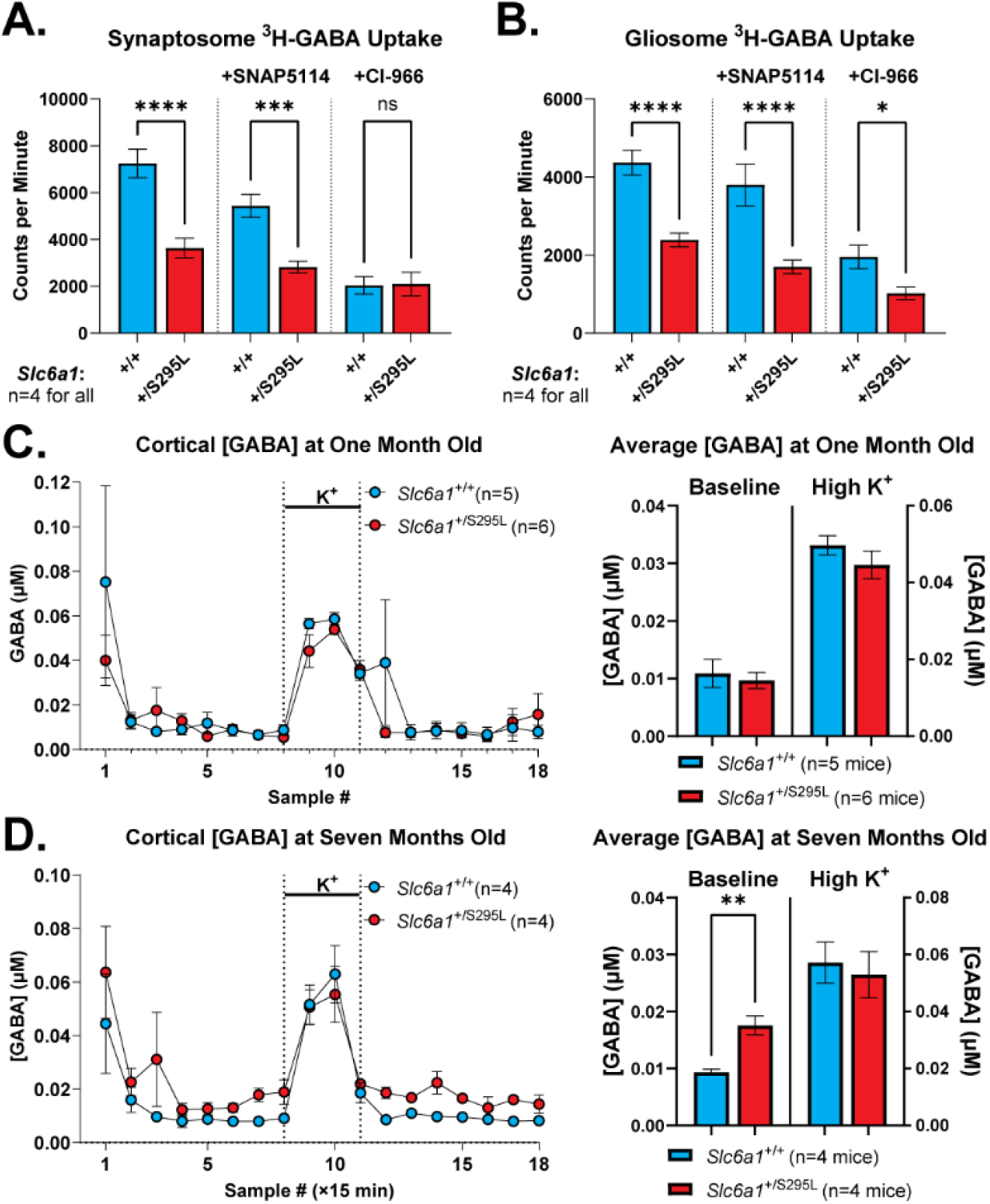
Deficient GABA uptake and elevation in extrasynaptic [GABA] in *Slc6a1*^+/S295L^ mice. **(A**,**B)** 3H-GABA uptake measured in synaptosomes **(A)** and gliosomes **(B)** extracted from brains of *Slc6a1*^+/S295L^ and *Slc6a1*^+/+^ littermates. Left to right, bar graphs show total uptake, GAT-1-dominant fraction in presence of GAT-3 blocker SNAP5114, and GAT-3-dominant fraction in presence of GAT-1 blocker Cl-966. **(C**,**D)** [GABA] measured by microdialysis in prefrontal cortex of *Slc6a1*^+/S295L^ and wildtype *Slc6a1*^+/+^ littermates at one month old **(C)** and seven months old **(D)** with heightened GABA release evoked with high K^+^ washon as indicated. Data are shown as means with SEM, ****, ***,**, *P<.0001,.001,.01,.05 by t-test.

We then determined if deficient uptake led to accumulation of extracellular GABA. We longitudinally measured [GABA] in prefrontal cortex in *Slc6a1*^+/S295L^ mice by microdialysis in vivo. We found an elevation in [GABA] with older age. At one month (Figure 1C), [GABA] was comparable in *Slc6a1*^+/S295L^ and *Slc6a1*^+/+^ sibling mice at baseline (.010 vs.011 µM WT) and during a spike in [GABA] elicited by high K^+^ washon (.045 vs.050 µM WT). However, at seven months (Figure 1D), [GABA] was significantly elevated in *Slc6a1*^+/S295L^ mice at baseline (.018 vs.0094 µM WT, **P<.01) and rose to levels comparable to wildtype during the K^+^-triggered spike (.053 vs.057 µM WT). A similar baseline increase in [GABA] has been previously reported in *Slc6a1* knockout mice together with an increased K^+^-triggered spike^12^. Elevated extracellular [GABA] suggests an elevation in GABA_A_ tonic currents, previously reported in *Slc6a1* knockout^12-15^. However, we did not detect a change in tonic GABA_A_ currents in cortex of 2-3 and 7 months old, and thalamus of one month old *Slc6a1*^+/S295L^ mice (Extended Data Figure 1), suggesting the existence of mechanisms for seizure generation other than increased tonic current. This also suggests that in these models, GABA_A_ tonic neurotransmission is not affected to the same extent as with complete knockout of *Slc6a1*, though it may still be elevated at physiological conditions *in vivo*.

We then evaluated synaptic transmission mediated by GABA_A_ receptors and found that synaptic responses in *Slc6a1*^+/S295L^ and *Slc6a1*^S295L/S295L^ mice were significantly prolonged due to deficient GAT-1 function (Figure 2). We recorded evoked and spontaneous inhibitory postsynaptic currents (eIPSCs and sIPSCs) in patch-clamped pyramidal neurons in acute cortical slices from *Slc6a1*^+/S295L^ and *Slc6a1*^S295L/S295L^ mice, and wildtype *Slc6a1*^+/+^ littermates. Compared to wildtype siblings, *Slc6a1*^+/S295L^ mice had prolonged eIPSCs (Figure 2A; median decay 57.5 vs 34.3 ms WT, **P<.01) and sIPSCs (Figure 2B; median decay 5.5 vs 4.1 ms WT, **P<.01). Similarly, *Slc6a1*^S295L/S295L^ mice had prolonged eIPSCs (Extended Data Figure 2; weighted tau 42.4 vs 18.4 ms WT, **P<.01) and sIPSCs (Figure 2C; mean decay 14.7 vs 10.4 ms WT, *P<.05). Duration of responses in wildtype mice significantly depended on GAT-1 activity, since application of GAT-1 blocker tiagabine prolonged eIPSCs greatly (Figure 2A, 110% decay increase, ***P<.001) and sIPSCs moderately (Figure 2B, 11% decay increase, ***P<.001). In comparison, GAT-1 function was greatly reduced in *Slc6a1*^+/S295L^ mice, where tiagabine affected eIPSCs significantly less (Figure 2A, 34% increase, *P<.05; **P<.01 vs wildtype change) and did not affect sIPSCs (Figure 2B, *P<.05 vs wildtype change). To test whether diminished GABA uptake had presynaptic effects, we also recorded three eIPSCs in quick succession and calculated the pulse ratios (Extended Data Figure 3). While amplitudes of successive eIPSCs diminished, there was no difference between genotypes or tiagabine treatment. However, we again saw significant prolongation of eIPSCs that led to event summation in case of successive events. In all experiments, eIPSCs appeared more sensitive to GAT-1 loss than sIPSCs.

**Figure 2.**
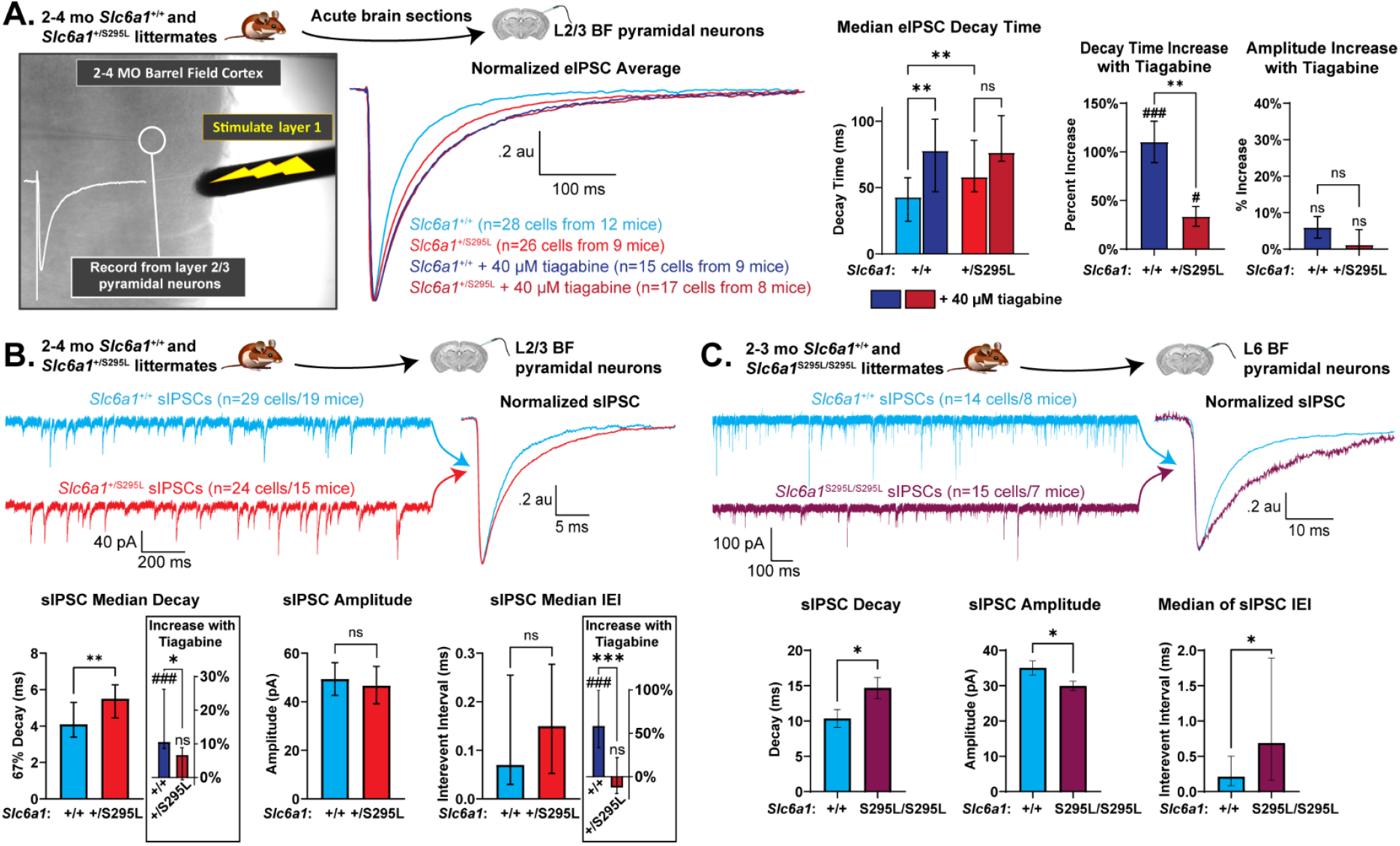
*Slc6a1*^+/S295L^ and *Slc6a1*^S295L/S295L^ mice have prolonged synaptic GABAergic currents and reduced sensitivity to GAT-1 block. **(A)** eIPSCs recorded from *Slc6a1*^+/S295L^ and *Slc6a1*^+/+^ littermates with and without GAT-1 blocker tiagabine. **(B, C)** sIPSCs recorded from *Slc6a1*^+/S295L^ **(B)** and *Slc6a1*^S295L/S295L^ **(C)** mice compared to *Slc6a1*^+/+^ littermates. Note that experiments in **(C)** are in L6 and at room temperature, compared to L2/3 at 32° C for **(A**,**B)**. eIPSC traces are normalized across-cell average, sIPSC traces and averaged sIPSC are from representative cells. Bar graphs represent means with SEM with t-tests or ANOVA with Sidak’s posthoc except where medians are noted, which show IQR with Kruskal-Wallis test with Dunn’s multiple comparisons. ***,**,*P<.001,.01,.05. For amplitude/decay increase with tiagabine in **(A-B)**, perneuron paired comparisons t-test was performed for each genotype. ###, #P<.0005,.025. Abbreviations: GAT-1, GABA transporter 1; mo, months old; BF, barrel field cortex; L, cortical layer; IEI, interevent interval; sIPSCs and eIPSCs, spontaneous and evoked inhibitory postsynaptic currents.

Likely, this is because the larger eIPSCs rely to greater extent on active GAT-1-mediated GABA clearance, while passive diffusion-mediated clearance plays a greater role for sIPSCs. These observations are in line with prior studies, which similarly show robust prolongation of eIPSCs when GAT-1 is knocked out or blocked, though only one study found such changes in sIPSCs^11,12,14,15^. In addition to changes in event duration, sIPSC in *Slc6a1*^S295L/S295L^ mice showed decreased amplitude (Figure 2C; mean 29.9 vs 35.0 pA WT, *P<.05) and increased interevent interval (Figure 2C; median.69 vs.21 ms WT, *P<.05), but not in *Slc6a1*^+/S295L^ mice (Figure 2B). These changes are likely specific to compromised GAT-1 function, and we also saw an increase in sIPSC interevent interval with tiagabine washon in wildtype mice (Figure 2B, 46% increase, ***P<.001).

We then evaluated if PBA can restore GABA uptake and synaptic transmission in *Slc6a1*^+/S295L^ mice. Acute 7-day PBA treatment significantly increased GABA uptake in *Slc6a1*^+/S295L^ synaptosomes (Figure 3A; 6057 vs 3641 CPM untreated, **P<.01) and gliosomes (Figure 3B; 3555 vs 2386 CPM untreated, *P<.05). This increase appeared specific to restoring GAT-1 function in *Slc6a1*^+/S295L^ mice, as PBA treatment restored GAT-1-mediated uptake observed with SNAP5114 in *Slc6a1*^+/S295L^ synaptosomes (Figure 3A; 4766 vs 2825 CPM untreated, **P<.01) and gliosomes (Figure 3B; 2892 vs 1698 CPM untreated,***P<.001), but did not affect uptake in wildtype synaptosomes or gliosomes (Extended Data Figure 4), and did not affect GAT-1-independent uptake with CL-966 in *Slc6a1*^+/S295L^ synaptosomes (Figure 3A; 1950 vs 2098 CPM untreated, ns) and gliosomes (Figure 3B; 1300 vs 1023 CPM untreated, ns).

**Figure 3.**
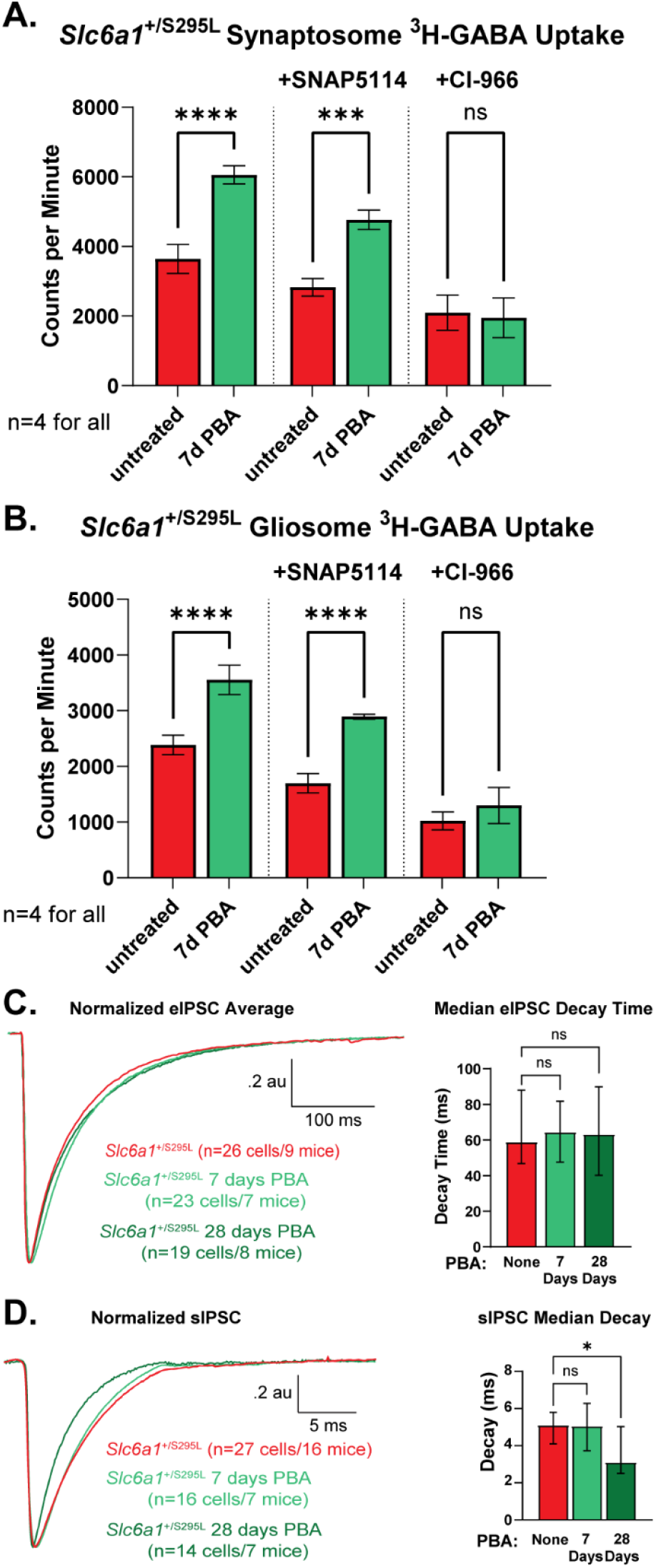
4-phenylbutyrate (PBA) treatment rescues GABA uptake and partially restores synaptic transmission in *Slc6a1*^+/S295L^ mice. **(A**,**B)** 3H-GABA uptake measured in synaptosomes **(A)** and gliosomes **(B)** extracted from brains of *Slc6a1*^+/S295L^ mice without treatment or after acute 7-day PBA treatment. Left to right, bar graphs show total uptake, GAT-1-dominant fraction in presence of GAT-3 blocker SNAP5114, and GAT-3-dominant fraction in presence of GAT-1 blocker Cl-966. **(C**,**D)** Average eIPSC traces (C) and representative sIPSC traces **(D)** recorded from *Slc6a1*^+/S295L^ mice without treatment, after acute 7-day i.p. or chronic oral 28-day treatment with 100 mg/kg/day PBA, with decay time quantified in bar graphs. Recording conditions were L2/3 pyramidal neurons in barrel field cortex with L1 stimulation at 32° C. Untreated *Slc6a1*^+/S295L^ data in **(A-D)** includes the data from *Slc6a1*^+/S295L^ mice in Figures 1 and 2. Data are shown as: **(A**,**B)** means with SEM with t-tests; **(C, D)** medians with IQR with Kruskal-Wallis test with Dunn’s multiple comparisons. ****,***,*P<.0001,.001,.05.

PBA treatment also restored duration of sIPSC in *Slc6a1*^+/S295L^ mice but not eIPSCs, and this required a longer treatment than restoring GABA uptake or seizures in prior studies^18,23^. We treated *Slc6a1*^+/S295L^ mice with one of two PBA regimens: acute intraperitoneal 7-day 100 mg/kg/day PBA or chronic oral 28-day 100 mg/kg/day PBA. *Slc6a1*^+/S295L^ mice receiving the chronic treatment had shorter sIPSCs (Figure 3D; median decay 3.1 vs 5.1 ms untreated; *P<.05), though eIPSCs remained prolonged (Figure 3C; median decay 63.2 vs 59.0 untreated). The acute treatment did not restore sIPSCs (Figure 3D; median decay 5.1 vs 5.1 untreated) nor eIPSCs (Figure 3C; median decay 64.4 vs 59.0 untreated). To address hypotheses in the field regarding PBA directly modulating GAT-1 function, we also washed on 10 mM PBA onto the slices during the recording, but found little effect on eIPSC or sIPSC duration or amplitude (Extended Data Figure 5).

Building upon prior work, present findings further define the pathomechanism of *SLC6A1*-DEEs (Extended Data Figure 6) and demonstrate PBA’s potential as a mechanism-based therapy. Previous studies of GABAergic neurotransmission focused on complete GAT-1 loss^11,12,14,15^, and we provide the first evidence in a patient-relevant model of a deficit of GABA clearance that leads to accumulation of GABA levels in the brain and pathologic changes in synaptic transmission. Our findings also highlight the therapeutic potential of PBA. Previously, we have shown that PBA can rescue the molecular pathology, seizures, and locomotor behaviors ^18,22,25^. We here demonstrate that impaired GABA uptake results in chronic extracellular GABA accumulation and impaired GABAergic neurotransmission, and that these abnormalities are partially rescued by PBA. Together, these findings provide compelling evidence that although PBA may act through multiple mechanisms, its therapeutic effect in *SLC6A1* variant–mediated DEEs involves restoration of extracellular GABA homeostasis and GABAergic neurotransmission through recovery of GAT-1-mediated uptake.

## Acknowledgements

We would like to acknowledge Dr. Mohammad Khan for help with microdialysis experiments. The work was carried out at Vanderbilt University Medical Center and was supported by research grant of National Institute of Health (NINDS) NS121718 and SLC6A1 Connect to JQK. GABA brain microdialysis experiments were partly supported by UCB. KZ postdoctoral training was supported by American Epilepsy Society Postdoctoral Research Fellowship and Vanderbilt Postdoctoral Training Program in Functional Neurogenomics (T32 NIMH).

## Author Contributions

Study design: KZ and JQK; Electrophysiology: KZ; Microdialysis: KR, MB, JQK; synaptosomal/gliosomal GABA uptake: WS, JQK; Animal husbandry and regulatory compliance: KR; Data analysis: KZ, KR, MB; Funding Acquisition: JQK; Supervision: JQK; Figure preparation and final data interpretation: KZ; KJQ; Writing: KZ and JQK. All authors reviewed and approved the manuscript.

