## Supplementary material for "Accumulation of Extracellular GABA, Impaired GABAergic Neurotransmission and 4-Phenylbutyrate Rescue in Mice of *SLC6A1* Variant-Mediated Disorders": Online Methods

### Mice Used in Experiments

The *Slc6a1*$^{+/S295L}$ mouse was created using Crispr-CAS9 knock-in by Shanghai Model Organisms (Shanghai Model Organisms Center, Inc., Cat. No. NM-KI-190014) and maintained in C57BL/6J background (Jax Stock #000664). Both male and female *Slc6a1*$^{+/S295L}$ mice, wildtype siblings, and *Slc6a1*$^{S295L/S295L}$ mice were used in this study. Mouse age differed for experiments as specified. All procedures were approved by the Vanderbilt University Institutional Animal Care and Use Committee and were conducted in accordance with the NIH guide for the Care and Use of Laboratory Animals.

### Animal Husbandry

Non-breeder animals were housed with maximum capacity of five same-sex animals per cage. Cages were kept in a temperature-controlled room (22 ± 1°C) under a 12:12 h light/dark cycle, with food and water ad libitum. Animals were excluded from the study if they showed any visible distress, such as shallow breathing, decreased movement, excessive grooming, and scratching. Male and female *Slc6a1*$^{+/S295L}$ mice were used for breeding. Mice were weaned and tail snipped at P21, and genotyped by real-time PCR (forward 5'-GCCGCCACCCAGATCT-3'; reverse 5' TTGTAGCTTCCCAGAGCAATCAG-3') using Transnetyx Genotyping Services (Cordova, TN).

### PBA administration

PBA 100 mg/kg dose was based on our prior experiments treating seizures and behavioral co-morbidities in *Slc6a1*$^{+/S295L}$ mice[1] and toxicity experiments in wildtype mice[2], and is within comparable range with 12.4 g/m$^2$/day dosing used in human patients[3]. For acute treatment, mice were given 100 mg/kg PBA (Sigma-Aldrich, P21005; 20 mg/mL in 0.9% normal saline, neutralized with 1 M potassium hydroxide to pH 7.4; stored at 4°C) through daily intraperitoneal (i.p.) injections for 7 days. For chronic treatment, mice were administered 100 mg/kg PBA via oral feeding in 75 mg peanut butter (Jif, smooth) presented on a small wooden stick daily for 28 days, preceded by 3 days peanut butter training without PBA. Testing started on the last day at least 2 hr after the final treatment.

### Synaptosome preparation

Synaptosome fractions were prepared as previously described[4]. All steps were performed at 4°C using pre-chilled buffers containing protease inhibitors. Brains were rapidly removed (<1 min after decapitation), and regions of interest were dissected and homogenized in HEPES-buffered sucrose, HBS (0.32 M sucrose, 10 mM HEPES, 2 mM EDTA, pH 7.4) at a ratio of 10 mL buffer per g tissue using a glass-Teflon homogenizer (10–15 strokes). Homogenates were centrifuged at 1,000 × g for 10 min, and the resulting supernatant was centrifuged at 13,800 × g for 20 min to obtain the crude synaptosomal pellet (P2). The pellet was resuspended in HBS (1 mL per 0.1 g tissue), recentrifuged under the same conditions, and kept on ice for subsequent uptake assays.

### Gliosome preparation

For gliosome isolation, homogenates were prepared in 0.32 M sucrose/25 mM Tris (pH 7.4), centrifuged sequentially at 1,000 × g for 5 min and 12,000 × g for 5 min, and the pellet was resuspended and loaded onto a Percoll gradient (20%, 10%, 6%, 2%) prepared in the same buffer. Gradients were centrifuged at 33,500 × g for 30 min, and the gliosome fraction was collected from the 2–6% Percoll interface, washed twice in PBS (20,000 × g for 10 min each), and resuspended in 25 mM HEPES (pH 7.4) for immediate use in $^3$H-GABA uptake experiments.

**GABA uptake**

GABA uptake was measured as previously described[4]. Synaptosome and gliosome fractions were prepared as described above and kept on ice until use. Each pellet was resuspended in 200 µL phosphate-buffered saline (PBS) and mixed with 1 mL preincubation buffer (in mM: 140 NaCl, 5 KCl, 1 $MgSO_4$, 2 glucose, 2.5 HEPES; pH 7.4, 310 mOsm) containing 10 µM unlabeled GABA and 5 µM [$^3$H]GABA (1 mCi/mL; PerkinElmer). Suspensions were incubated at room temperature (22–24 °C) for 30 min with gentle mixing to allow uptake. Uptake was terminated by rapid vacuum filtration through 20 µm nylon filters (Thermo Fisher) mounted on a 50 mm manifold. The filtrate was discarded, and filters retaining synaptosomes or gliosomes were washed three times with 1 mL ice-cold preincubation buffer to remove unbound tracer. Filters were then air-dried for 5 min at room temperature, transferred to 35 mm dishes containing 750 µL of 10% SDS, and gently agitated until the material was fully solubilized (5–10 min). Aliquots of the lysate (650 µL) were mixed with 5 mL scintillation cocktail (Bio-Safe II, RPI) and quantified for $^3$H β-emission using a liquid scintillation counter. For inhibitor conditions, samples were incubated with 50 µM CI-966 or 30 µM SNAP-5114 added to the uptake buffer. Uptake was expressed as counts per minute (CPM).

**Microdialysis**

*Slc6a1*$^{+/S295L}$ mice and wildtype siblings were anesthetized with 1.5% isoflurane and secured in a stereotaxic chamber with bregma-lambda < ± 0.1. The Microdialysis canula was planted in the frontal cortex with coordinates of 1.0 mm (Medial-lateraL), 2.0 mm (anterior-posterior) and 2.25 mm in depth. After 7 days recovery from the surgery, the mice were subjected for sample collection in Artificial cerebrospinal fluid (aCSF). A total of 18 fractions with a total volume of 30 µL for each fraction at a rate of 2 µL/min were collected. The 9th and 10th fractions were collected after switching to the aCSF with 50 mM KCl. The samples were collected on ice and stored at -80ºC.

GABA concentration in microdialysis samples was then quantified by the Vanderbilt University Neurochemistry Core using liquid chromatography/mass spectrometry (LC/MS) methodology following derivatization with benzoyl chloride (BZC). Briefly, 5 µL of sample fluid was added directly to a 300 µL Verex LC/MS vial (Phenomenex, Torrance, CA USA). 10 µL each of 500 mM NaCO3 (aq) and 2% BZC in acetonitrile was added to the vial and vortexed. After two minutes, the reaction was stopped by the addition of 10 µL internal standard solution. After this, LC/MS was performed on a 2.0 x 50 mm, 1.7 mm particle Acquity BEH C18 column (Waters Corporation, Milford, MA, USA) using a Waters Acquity UPLC. Mobile phase A was 15% aqueous formic acid, and mobile phase B was acetonitrile. Samples were separated by a gradient of 98–5% of mobile phase A over 11 min at a flow rate of 600 mL/min prior to delivery to a Waters Xevo TQ-XS triple quadrupole mass spectrometer.

**Preparation of Acute Mouse Brain Slices for Recording**

Slicing protocol was based on Ting et al, 2018[5]. Mice were anesthetized with isoflurane and transcardially perfused at 3.5-5 mL/min with cold, oxygenated NMDG cutting solution, consisting of (mM): 92 NMDG, 2.5 KCl, 1.25 $NaH_2PO_4$, 30 $NaHCO_3$, 20 HEPES, 25 glucose, 2 thiourea, 5 Na-ascorbate, 3 Na-pyruvate, 0.5 $CaCl_2 \cdot 2H_2O$, and 10 $MgSO_4 \cdot 7H_2O$ (300-305 mOsm, titrated to pH 7.3–7.4 with 5 M hydrochloric acid). For thalamic recordings, sucrose cutting solution was used, which consisted of (mM): 200 sucrose, 1.9 KCl, 1.2 $NaH_2PO_4$, 6 $MgCl_2$, .5 $CaCl_2$, 10 glucose, 25 $NaHCO_3$ (305 mOsm, pH 7.4). Mice were then decapitated, their brains were dissected, blocked, and coronal slices were sectioned at 300 µm on a Leica VT1200S vibratome at a speed of .06 mm/s in cold, oxygenated cutting solution. Slices were then recovered for 30 min at 34° C in oxygenated holding solution, consisting of (mM): 92 NaCl, 2.5 KCl, 1.25 $NaH_2PO_4$, 30 $NaHCO_3$, 20 HEPES, 25 glucose, 2 thiourea, 5 Na-ascorbate, 3 Na-pyruvate, 2

$CaCl_2 \cdot 2H_2O$, and 2 $MgSO_4 \cdot 7H_2O$ (300-305 mOsm, pH 7.4), except thalamic recordings, where artificial cerebrospinal fluid (aCSF) was used instead. Subsequently, slices were recovered for 1 hr at 20-21°C in oxygenated aCSF, consisting of (mM): 124 NaCl, 2.5 KCl, 1.25 $NaH_2PO_4$, 24 $NaHCO_3$, 12.5 glucose, 5 HEPES, 2 $CaCl_2 \cdot 2H_2O$, and 2 $MgSO_4 \cdot 7H_2O$ (300-305 mOsm, pH 7.4).

**Mouse Acute Brain Slice Recording and Analysis**

Recordings were performed in a chamber perfused with oxygenated aCSF at a rate of 2.0 mL/min. Neurons were patched with 3-5 MΩ borosilicate glass electrodes (World Precision Instruments, TW150-4, pulled with Narishige PP-830 or Sutter P-2000 pullers) filled with internal solution containing (mM): 150 CsCl, 1 $MgCl_2$, 10 HEPES, .1 $CaCl_2$, 1.1 EGTA, and 2 $Na_2ATP$ (285-290 mOsm, pH 7.4) and clamped at -70 mV. Whole-cell recordings were acquired using an Axon MultiClamp 700B amplifier, filtered at 2 kHz, digitized at 10 kHz with Digidata 1440A, and recorded with Clampex 10.4 software (Molecular Devices).

Recordings in ventrobasal thalamic neurons and somatosensory cortex layer 6 pyramidal neurons (tonic current, sIPSCs) were done at 20-21°C in presence 10 μM NBQX and 50 μM APV, while recordings in layer 2/3 somatosensory cortex pyramidal neurons (eIPSCs, sIPSCs) were done at 32°C in presence 10 μM NBQX, 50 μM APV, and 500 nM CGP-54626 with addition of 40 μM tiagabine or 10 mM PBA as noted. Somatosensory cortex was identified in reference to hippocampus and apparent barrels in barrel field cortex. Cortical layers and pyramidal neurons were identified based on morphology under DIC. Ventrobasal thalamus (VB) was identified as a darkened, almond-shaped structure with a striated appearance adjacent to the internal capsule and below the hippocampus.

sIPSCs were recorded in aCSF and analyzed with EasyElectrophysiology software. For layer 6 sIPSCs, manual detection and 15 pA event threshold were used. For layer 2/3 large sIPSCs, automatic detection and event threshold set at 25 pA were used. Series resistance ($R_a$) was uncorrected, and recordings were discarded if $R_a$ was over 25 MΩ or changed by more than 15% over the course of a recording. Representative average sIPSCs were calculated using EasyElectrophysiology. Tonic current was recorded by bath-application of 30 μM bicuculline in addition to APV and NBQX, which caused a baseline shift and blocked all events. Using ClampFit software (Molecular Devices), average baseline over 30 s period was calculated (A) a few minutes before, (B) immediately before, and (C) after the baseline shift. To exclude events from analysis, baseline average was calculated by fitting a curve to the right half of all-points histogram, and measuring the peak value. The difference (C)-(B) was used to calculate the tonic current, and the difference (B)-(A) was used to control for stability of the baseline during the recording.

For eIPSCs, a stimulating electrode (FHC 30200) was placed in layer 1 of somatosensory cortex. 200-800 μA of stimulation was applied for 100 μs to evoke a single 300-900 pA event in a patched cell. Series resistance ($R_a$) was monitored same as for other recordings. eIPSCs were analyzed by creating an average event for each cell in Clampfit, after which the average events for each cell were fitted and analyzed in EasyElectrophysiology, from which average amplitude, decay, rise time, and area under the curve were calculated, and an average trace per condition was made. eIPSCs with unusually slow kinetics were discarded from analysis.

**Data analysis and statistics**

Statistical data was stored using Microsoft Excel, analyzed and graphed using GraphPad Prism. Final figures were made in Adobe Illustrator. Statistical tests are specified in each figure. Values are displayed as mean ± SEM with t-test or one-way ANOVA with Sidak's posthoc test except where normality test was not passed, which are displayed as median ± IQR with Mann-Whitney or Kruskal-Wallis test with Dunn's posthoc test. Paired-comparisons t-test or Dunn's paired comparisons test were used for change with washon experiments. ****, ***, **, * P<.0001, .001, .01, .05, respectively.
