## Extended Data Figures for "Accumulation of Extracellular GABA, Impaired GABAergic Neurotransmission and 4-Phenylbutyrate Rescue in Mice of *SLC6A1* Variant-Mediated Disorders"

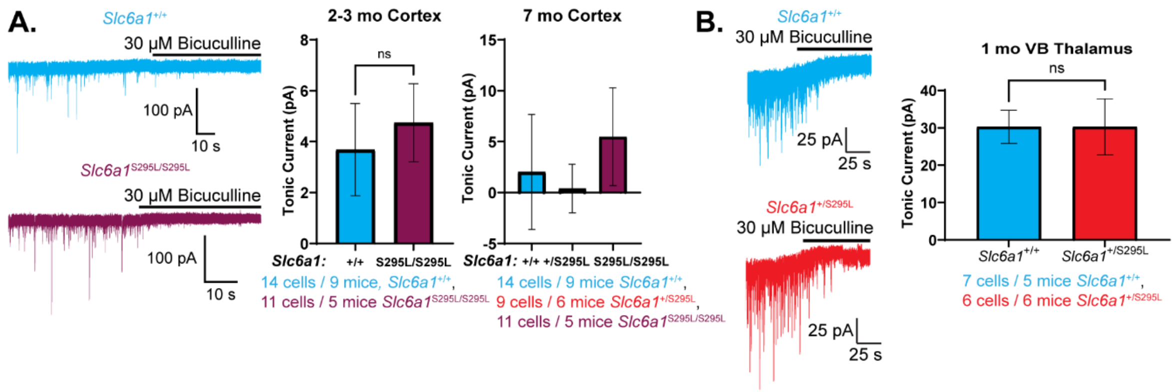


**Extended Data Figure 1. No change in tonic GABA$_A$ receptor-mediated current in cortex and thalamus of *Slc6a1*$^{S295L/S295L}$ and *Slc6a1*$^{+/S295L}$ mice, respectively. A.** Representative traces of tonic current observed with washon of GABA$_A$ receptor blocker bicuculline in layer 6 somatosensory cortex pyramidal neurons of *Slc6a1*$^{S295L/S295L}$ and wildtype *Slc6a1*$^{+/+}$ littermates at 2-3 and 7 months old with quantification on the right. **B.** Representative traces of tonic current observed with washon of bicuculline in ventrobasal thalamus neurons of *Slc6a1*$^{+/S295L}$ and wildtype *Slc6a1*$^{+/+}$ littermates at 1 month old with quantification on the right. **A, B.** Recordings done at room temperature. Graphs show mean with SEM with t-tests or ANOVA with Sidak's posthoc test.

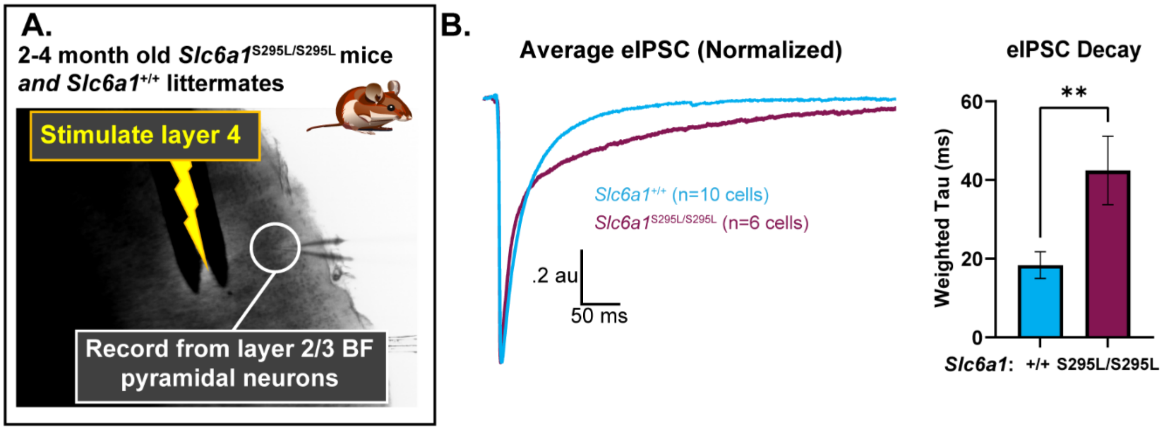


**Extended Data Figure 2. *Slc6a1*$^{S295L/S295L}$ mice show prolonged evoked inhibitory postsynaptic currents (eIPSCs). A.** Setup for recording eIPSCs in barrel field (BF). **B.** Average normalized eIPSC trace for *Slc6a1*$^{S295L/S295L}$ and wildtype *Slc6a1*$^{+/+}$ littermates with decay quantification on the right. Weighted tau was calculated for each cell after fitting a 5-10 trace average with 2-3 term exponential. Graph shows mean with SEM; **P<.01 by t-test.

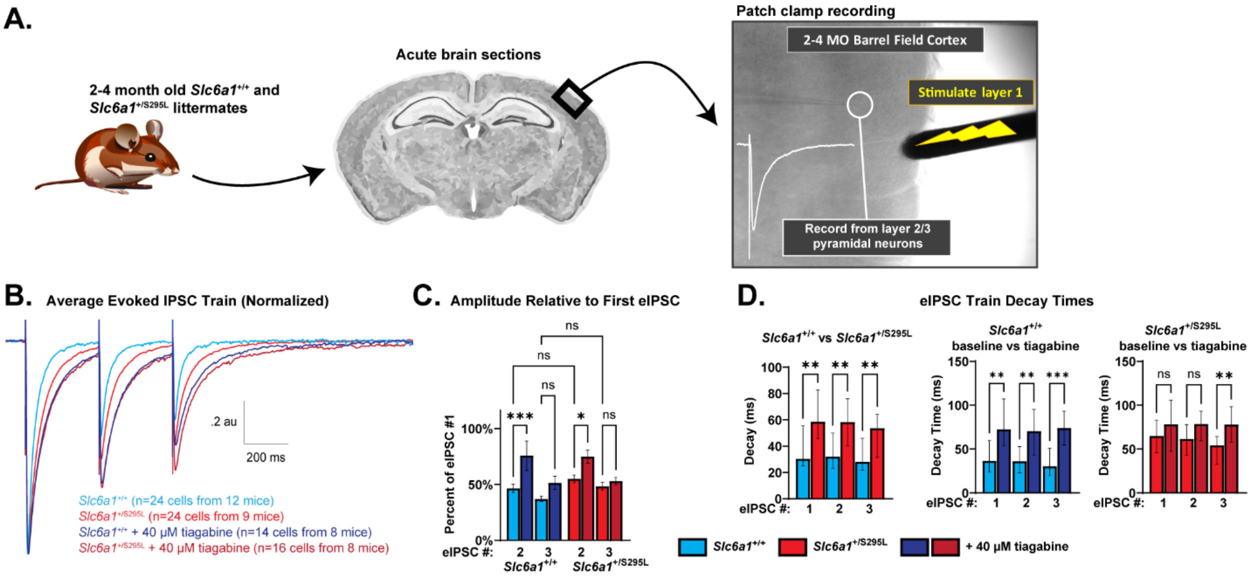


**Extended Data Figure 3. Prolonged duration and comparable amplitude depression in successive train-eIPSCs in *Slc6a1*$^{+/S295L}$ mice. A.** Experimental workflow for recording train-eIPSCs in response to .33 Hz 1 s stimulation in *Slc6a1*$^{+/S295L}$ and wildtype *Slc6a1*$^{+/+}$ littermates. **B.** Averaged train-eIPSC traces normalized on the first eIPSC. **C.** Amplitudes of second and third eIPSCs as percentage of the first eIPSC amplitude, shown as means ± SEM with ANOVA with Sidak's posthoc. **D.** Train-eIPSC decay shown as medians ± IQR with Kruskal-Wallis (baseline) or Dunn's paired comparisons (tiagabine) test.

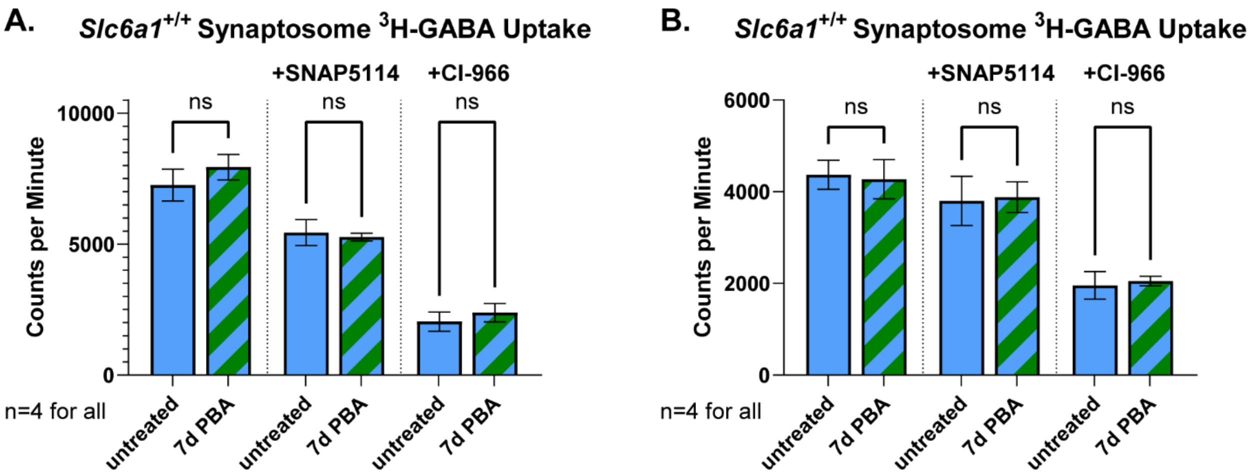


**Extended Data Figure 4. 4-phenylbutyrate (PBA) treatment does not affect GABA uptake in wildtype *Slc6a1*$^{+/+}$ mice. (A,B)** 3H-GABA uptake measured in synaptosomes **(A)** and gliosomes **(B)** extracted from brains of wildtype *Slc6a1*$^{+/+}$ mice without treatment or after acute 7-day PBA treatment. Left to right, bar graphs show total uptake, GAT-1-dominant fraction in presence of GAT-3 blocker SNAP5114, and GAT-3-dominant fraction in presence of GAT-1 blocker CI-966. Untreated *Slc6a1*$^{+/+}$ data in **(A-D)** includes the data from *Slc6a1*$^{+/+}$ mice in Figures 1. Data are shown as means ± SEM with t-tests.

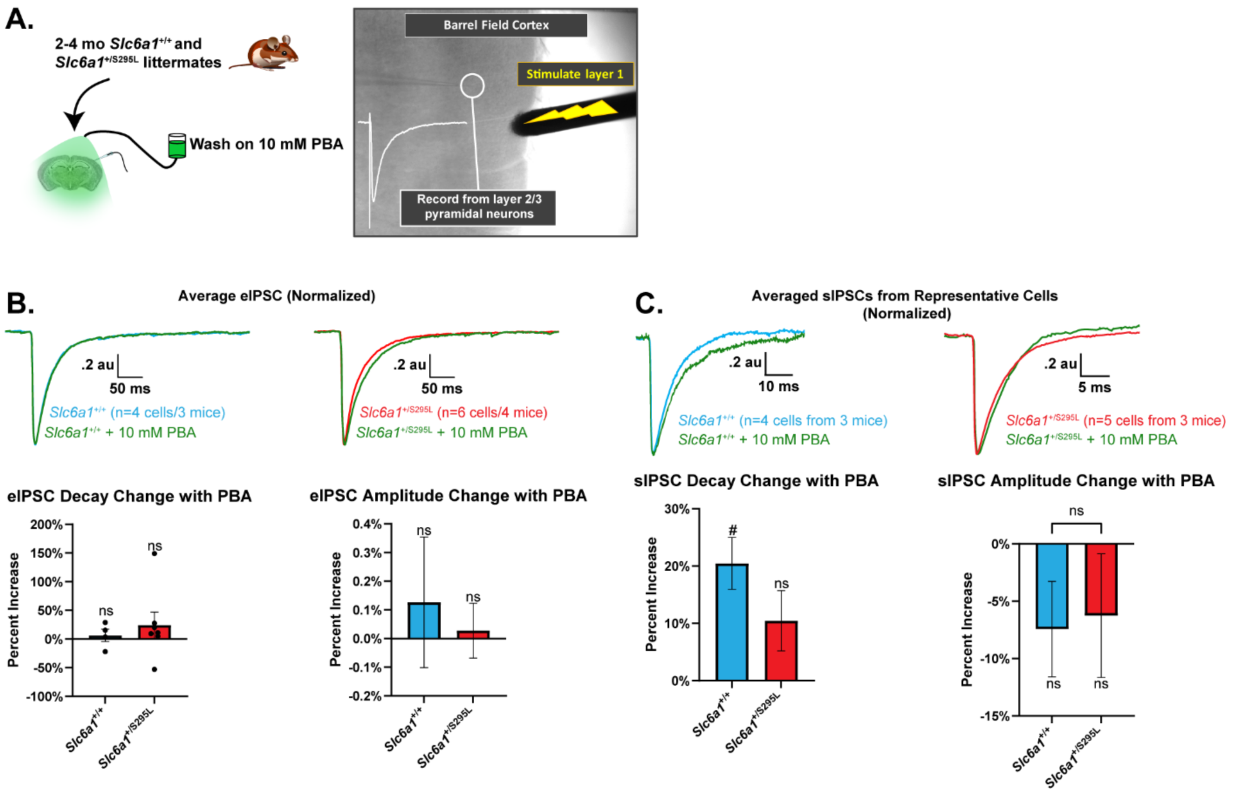


**Extended Data Figure 5. Direct PBA application does not restore duration of eIPSCs and sIPSC in *Slc6a1*$^{+/S295L}$ mice. A.** Experimental workflow for recording eIPSCs; 10 mM PBA was washed on after an eIPSC baseline was established. **B.** Average normalized eIPSC traces shown separately for wildtype *Slc6a1*$^{+/+}$ littermates and *Slc6a1*$^{+/S295L}$ mice before and after 10 mM PBA application with per-cell quantification of decay and amplitude change shown in bar graphs. **C.** Representative normalized sIPSC traces shown separately for wildtype *Slc6a1*$^{+/+}$ littermates and *Slc6a1*$^{+/S295L}$ mice before and after 10 mM PBA application with per-cell quantification of decay and amplitude change shown in bar graphs. **B,C.** Recordings were done at 32$^{O}$ C. Bar graphs show mean ± SEM with paired t-test for each condition (#P<.05) and a t-test between conditions.

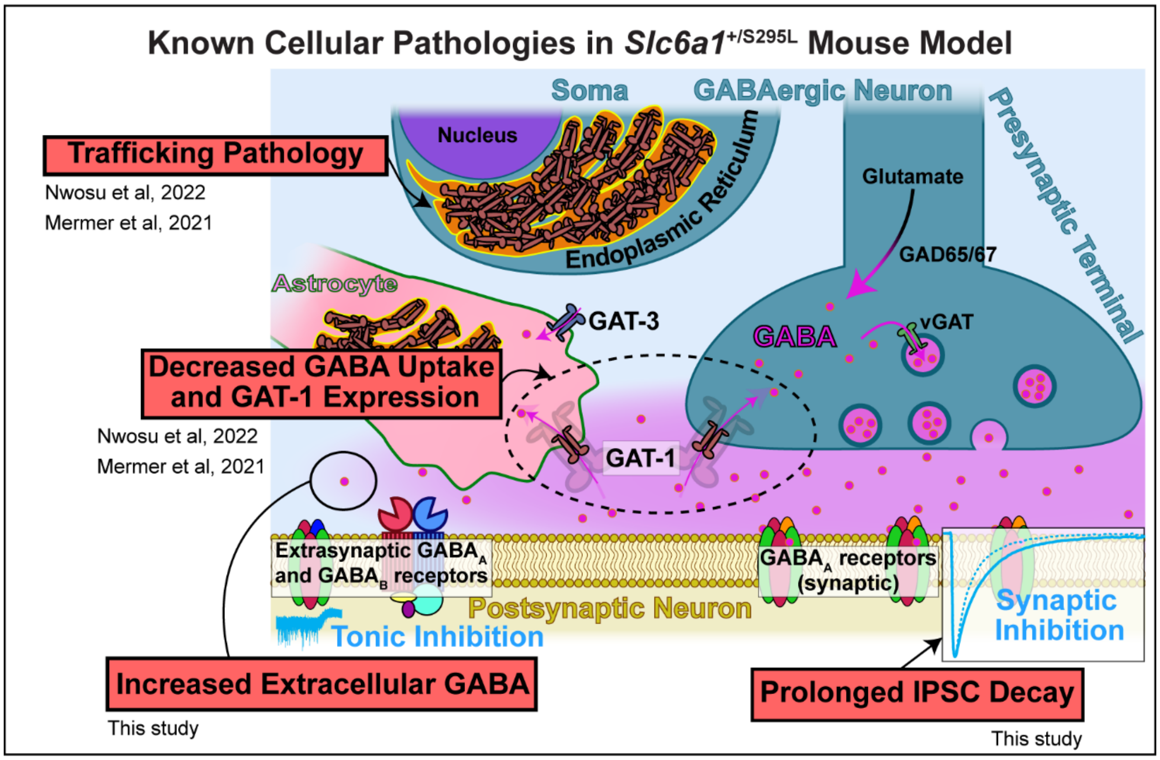


**Extended Data Figure 6. Summary of known pathologies in *SLC6A1*-DEE based on our earlier and current *Slc6a1*$^{+/S295L}$ mouse model studies.**
